# An African origin for *Mycobacterium bovis*

**DOI:** 10.1101/773192

**Authors:** Chloé Loiseau, Fabrizio Menardo, Abraham Aseffa, Elena Hailu, Balako Gumi, Gobena Ameni, Stefan Berg, Leen Rigouts, Suelee Robbe-Austerman, Jakob Zinsstag, Sebastien Gagneux, Daniela Brites

## Abstract

**Background and objectives:** *Mycobacterium bovis* and *Mycobacterium caprae* are two of the most important agents of tuberculosis (TB) in livestock and the most important causes of zoonotic TB in humans. However, little is known about the global population structure, phylogeography and evolutionary history of these pathogens.

**Methodology:** We compiled a global collection of 3364 whole-genome sequences from *M. bovis* and *M. caprae* originating from 35 countries and inferred their phylogenetic relationships, geographic origins and age.

**Results:** Our results resolved the phylogenetic relationship among the four previously defined clonal complexes of *M. bovis*, and another eight newly described here. Our phylogeographic analysis showed that *M. bovis* likely originated in East Africa. While some groups remained restricted to East- and West Africa, others have subsequently dispersed to different parts of the world.

**Conclusions and implications:** Our results allow a better understanding of the global population structure of *M. bovis* and its evolutionary history. This knowledge can be used to define better molecular markers for epidemiological investigations of *M. bovis* in settings where whole genome sequencing cannot easily be implemented.

## BACKGROUND AND OBJECTIVES

Tuberculosis (TB) remains an important burden for global health and the economy [1]. TB is the number one cause of human death due to infection globally, with an estimated 10.0 million new cases and 1.5 million deaths occurring every year [1]. TB is caused by members of the *Mycobacterium tuberculosis* complex (MTBC), which includes seven human-adapted lineages, and several animal-adapted ecotypes including *M. bovis* and *M. caprae.* Animal TB complicates the control of human TB due to the zoonotic transfer of TB bacilli from infected animals to exposed human populations e.g. through the consumption of unpasteurized milk or handling of contaminated meat [2]. *M. bovis* and *M. caprae* are the most important agents of TB in livestock and the most important agents of zoonotic TB in humans, causing an estimated 147 000 new human cases and 12 500 human deaths yearly [1, 3]. Zoonotic TB caused by *M. bovis* also poses a challenge for patient treatment, due to its natural resistance to pyrazinamide (PZA), one of the four first-line drugs used in the treatment of TB. In addition, TB in livestock accounts for an estimated loss of three billion US dollars per year [4]. In Africa, the prevalence of *M. bovis* is highest in peri-urban dairy belts of larger cities and remains at low levels in rural areas [5], often also threatening wildlife populations [6]. During the last few years, analyses of large globally representative collections of whole genome sequences (WGS) from the human-adapted MTBC lineages have enhanced our understanding of the global population structure, phylogeography and evolutionary history of these pathogens [7]. By contrast, little corresponding data exist for the various animal-adapted ecotypes of the MTBC such as *M. bovis*.

Current knowledge about global *M. bovis* populations stems mostly from spoligotyping [8, 9]. This method has been highly valuable for showing that *M. bovis* populations vary by geography, and defining strain families based on the presence or absence of spacers in the Direct Repeat region of the MTBC genome [8]. However, the discriminatory capacity of spoligotyping is limited since diversity is measured at a single locus prone to convergent evolution and phylogenetic distances cannot be reliably inferred [10].

In addition to spoligotyping, other genomic markers such as deletions [11–14] and single nucleotide polymorphisms (SNPs) [14], have given insights into the biogeography of *M. bovis*. These markers have been used to define four major groups of genotypes within *M. bovis*, known as clonal complexes European 1 and 2 (Eu1, Eu2) and African 1 and 2 (Af1 and Af2) [11–14]. Bovine TB in West Africa and East Africa is mainly caused by the clonal complexes Af1 and Af2, respectively [11, 12]. Bovine TB in Europe and in the Americas is caused by clonal complex Eu1, which affects mostly the British Islands and former trading countries of the UK [13], while Eu2 is prevalent mostly in the Iberian Peninsula and Brazil [14].

More recently, studies based on WGS have brought deeper insights into the population dynamics of *M. bovis* and showed that unlike *M. tuberculosis*, wild animals can act as *M. bovis* reservoirs in different regions of the world [15–18]. However, most studies using WGS have aimed at investigating local epidemics, and little is known about the global population structure and evolutionary history of *M. bovis*. Recently, we suggested a scenario for the evolution of the animal-adapted MTBC, in which we propose that *M. caprae* and *M. bovis* might have originally come out of Africa [19]. Here we gathered 3356 *M. bovis* and *M. caprae* WGS from the public domain, to which we added eight new *M. bovis* sequences from strains isolated in East Africa. Our results provide a phylogenetic basis to better understand the global population structure of *M. bovis*. Moreover, they point to East Africa as the most likely origin of contemporary *M. bovis*.

## METHODS

### Data collection

A total of 3929 *M. bovis* genomes were retrieved from EBI: 3834 BioSamples were registered on EBI with the taxon id 1775 (corresponding to “*Mycobacterium tuberculosis* variant bovis”) and downloaded on the 11^th^ of March 2019 and 95 *M. bovis* genomes were registered under taxon id 1765 (corresponding to “*Mycobacterium tuberculosis”*).

Of these, 457 were excluded because they were part of pre-publications releases from the Wellcome Trust Sanger Institute, 130 were excluded because they were registered as BCG – Bacille Calmette Guérin, the vaccine strain derived from *M. bovis*, one genome was excluded because it was wrongly classified as *M. bovis*, and three samples were excluded because they corresponded to RNA-seq libraries.

In addition, we added 81 publically available *M. caprae* genomes and eight previously unpublished sequences from *M. bovis* isolated in Ethiopia (n=7) and Burundi (n=1). The sequencing data has been deposited in the European Nucleotide Archive (EMBL-EBI) under the study ID PRJEB33773.

From this total of 3427 genomes, 63 sequences were excluded because they did not meet our criteria for downstream analyses (average whole-genome coverage below 7, ratio of heterogeneous SNPs to fixed SNPs above 1), yielding a final dataset of 3364 genomes (Fig. S1, Table S1). Geographical origin of the isolates, date of isolation and host metadata were recovered from EBI (Table S1).

### Whole genome sequence analysis

All samples were subject to the same whole-genome sequencing analysis pipeline, as described in [20]. In brief, reads were trimmed with Trimmomatic v0.33 [21]. Only reads larger than 20 bp were kept for the downstream analysis. The software SeqPrep (https://github.com/jstjohn/SeqPrep) was used to identify and merge any overlapping paired-end reads. The resulting reads were aligned to the reconstructed ancestral sequence of the MTBC [22] using the MEM algorithm of BWA v0.7.13 [23] with default parameters. Duplicated reads were marked using the MarkDuplicates module of Picard v2.9.1 (https://github.com/broadinstitute/picard). The RealignerTargetCreator and IndelRealigner modules of GATK v 3.4.0 were used to perform local realignment of reads around InDels [24]. Finally, SNPs were called with Samtools v1.2 mpileup [25] and VarScan v2.4.1 [26] using the following thresholds: minimum mapping quality of 20, minimum base quality at a position of 20, minimum read depth at a position of 7X, maximum strand bias for a position 90%. Only SNPs considered to have reached fixation within an isolate (frequency within-isolate ≥90%) were considered. For SNPs with ≤10% frequency, the ancestor state was called. SNPs were annotated using snpEff v4.1144 [27], using the *M. tuberculosis* H37Rv reference annotation (NC_000962.3) as the genome of *M. bovis* (AF2122/97) has no genes absent from H37Rv except for TbD1 (contains *mmpS6* and the 5′ region of *mmpL6*) [28].

### *In silico* spoligotyping, genomic deletions and previously defined clonal complexes

The spoligotype pattern of the 3364 genomes was determined *in silico* using KvarQ [29]. The results were submitted to the *Mycobacterium bovis* spoligotype database https://www.mbovis.org/ [30] and SB numbers obtained.

All 3364 genomes were screened *in silico* for the presence of molecular markers defining the previously described *M. bovis* clonal complexes; i.e. for the presence or absence of the genomic deletions RDAf1, RDAf2, RDEu1 (also known as RD17) [11–13], and in the case of Eu2 [14], for the presence of SNP 3813236 G to A with respect to the H37Rv (NC_000962.3). Other deletions, RD4, RDpan and N-RD17, previously used to genotype *M. bovis* lineages were also screened for [31–33]. The genomic coordinates in H37Rv (NC_000962.3) used to determine each deletion were the following; RDAf1 (664254-669601); RDAf2 (680337-694429); RDEu1 (1768074-1768878); RD4 (1696017-1708748); RDpan (4371020-4373425); N-RD17 (3897069-3897783). A genomic region was considered deleted if the average coverage over the region was below two.

### Phylogenetic analyses

All phylogenetic trees were inferred with RAxML (v.8.2.12) using alignments containing only polymorphic sites. A position was considered polymorphic if at least one genome had a SNP at that position with a minimum percentage of reads supporting the call of 90%. Deletions and positions not called according to the minimum threshold of 7, were encoded as gaps. We excluded positions with more than 10% missing data, positions falling in PE/PPE genes, phages, insertion sequences and in regions with at least 50 bp identity to other regions in the genome [34]. Positions falling in drug resistance-related genes were also excluded. The alignment used to produce Figure 2 comprised 22 492 variable positions and the alignment used to produce Figure S2 comprised 45 981 variable positions.

Maximum likelihood phylogenies were computed using the general time-reversible model of sequence evolution (-m GTRCAT-V options), 1,000 rapid bootstrap inferences, followed by a thorough maximum-likelihood search performed through CIPRES [35]. All phylogenies were rooted using a *M. africanum* Lineage (L) 6 genome from Ghana (SAMEA3359865).

### Obtaining a representative dataset of *M. bovis* genomes - Subsampling 1

Our phylogenetic reconstruction indicated that sequences belonging to clonal complex Eu1 and Eu2 were over-represented in the initial 3364 genome dataset, particularly from the USA, Mexico, New Zealand and the UK. To obtain a smaller dataset with a more even representation of the different phylogenetic groups, we pruned the 3364 genomes using the following criteria: 1) we removed all genomes with non-available country metadata (n=739), which resulted in 2625 genomes; 2) we used Treemmer v0.2 [20] with the option *-RTL 99* to keep 99% of the original tree length and the option *–lm* to include a list of taxa to protect from pruning. This list included all genomes belonging to clonal complexes Af1 and Af2, as well as any genome belonging to any unclassified clade; 3) we visually identified monophyletic clades with all taxa from the same country and used Treemmer v0.2 [20], using options *-lmc* and *–lm*, to only keep a few representatives of each of these clades. To have representatives of the BCG clade, we kept 11 BCG genomes from [36]. This selection process rendered a dataset of 476 genomes.

### Ancestral reconstruction of geographic ranges - Subsampling 2

To infer the geographic origin of the ancestors of the main groups of *M. bovis* and *M. caprae*, we used the 476 genomes dataset (see subsampling 1) and excluded all BCG genomes and all *M. bovis* from human TB cases or from unknown hosts, if the strains were isolated in a low incidence TB country (Europe, North America, Oceania). This is justified by the fact that the majority of such cases correspond to immigrants from high incidence countries that were infected in their country of origin, i.e. country of isolation does not correspond to the native geographic range of the strain and is thus not informative for the geographic reconstruction. *M. bovis* from patients in high incidence countries were kept (Table S1). The resulting dataset was composed of 392 genomes.

For the ancestral reconstruction of geographic ranges, we used the geographic origin of the strains and the phylogenetic relationships of the 392 genomes. Geographic origin was treated as a discrete character to which 13 states, corresponding to UN-defined regions, were assigned. To select the best model of character evolution, the function fitMk from the package phytools 0.6.60 in R 3.5.0 [37] was used to obtain the likelihoods of the models ER (equal-rates), SYM (symmetrical) and ARD (all rates different) [38]. A Likelihood Ratio Test (LRT) was used to compare the different log-Likelihoods obtained. According to the former, the best fitting model was SYM, a model that allows states to transition at different rates in a reversible way, i.e. reverse and forward transitions share the same parameters (Table S2). The function *make.simmap* in phytools package 0.6.60 in R 3.5.0 [37, 39] was used to apply stochastic character mapping as implemented in SIMMAP [40] on the 392 genomes phylogeny inferred from the best-scoring ML tree rooted on L6, using the SYM model with 100 replicates. We summarized the results of the 100 replicates using the function *summary* in phytools package 0.6.60 in R [37].

### Molecular Dating of *M. bovis* and *M. caprae* – Subsampling 3

For the molecular clock analyses, we considered only genomes for which the date of isolation was known (n=2058). For the eight genomes sequenced in this study, the date of isolation was retrieved at a later point, and these strains were not included in the dating analysis (Table S1). We used a pipeline similar to that reported in [20]. We built SNP alignments including variable positions with less than 10% of missing data (alignment length; 24 828 variable positions). We added an L6 strain as outgroup (SAMEA3359865) and inferred the Maximum Likelihood tree as described above. Since the alignment contained only variable positions, we rescaled the branch lengths of the trees: rescaled_branch_length = ((branch_length * alignment_length) / (alignment_length + invariant_sites)). To evaluate the strength of the temporal signal, we performed root-to-tip regression using the R package ape [41]. Additionally, we used the least square method implemented in LSD v0.3-beta [42] to estimate the molecular clock rate in the observed data and performed a date randomization test with 100 randomized datasets. To do this, we used the quadratic programming dating (QPD) algorithm and calculated the confidence interval (options -f 100 and -s).

We also estimated the molecular clock rates using a Bayesian analysis. For this, we reduced the dataset to 300 strains with Treemmer v0.2 in the following way: we randomly subsampled strains, maintaining the outgroup and at least one representative of four small clades of the tree that would have disappeared with simple random subsampling strategy (Af2 clonal complex: G42133; Af1 clonal complex: G02538; *PZA_ sus_unknown1*: G04143, G04145, G04147; *M. caprae*: G42152, G42153, G37371, G37372, G41838; Table S1). The resulting alignment included 13 012 variable sites (subset1).

We used jModelTest 2.1.10 v20160303 [43] to identify the best fitting nucleotide substitution model among 11 possible schemes, including unequal nucleotide frequencies (total models = 22, options -s 11 and -f). We performed Bayesian inference with BEAST2 [44]. We corrected the xml file to specify the number of invariant sites as indicated here: https://groups.google.com/forum/#!topic/beast-users/QfBHMOqImFE, and used the tip sampling years to calibrate the molecular clock.

We used the uncorrelated lognormal relaxed clock model [45], the best fitting nucleotide substitution model according to the results of jModelTest (all criteria selected the transversional model (TVM) as the best model), and three different coalescent priors: constant population size, exponential population growth and the Bayesian Skyline [46]. We chose a 1/x prior for the population size [0-10^9^], a 1/x prior for the mean of the lognormal distribution of the clock rate [10^−10^ – 10^−5^], and the standard Gamma distribution as prior for the standard deviation of the lognormal distribution of the clock rate [0 – infinity]. For the exponential growth rate prior, we used the standard Laplace distribution [-infinity – infinity]. For all analyses, we ran two runs and used Tracer 1.7.1 [47] to evaluate convergence among runs and to calculate the estimated effective sample size (ESS). We stopped the runs when they reached convergence, and the ESS of the posterior and of all parameters were larger than 200. The number of generations ranged from 150 to 300 million depending on the run. We used Tracer [48] to identify and exclude the burn-in, which ranged from 4 to 30 million generations, depending on the run.

Since the BEAST analysis was based on a sub-sample of the data, we tested the robustness of the sub-sampling by repeating twice the sub-sampling with Treemmer [20]. This resulted in two alignments of 13 272 and 12 820 SNPs, respectively (subset2 and subset3). We then repeated all the BEAST analyses described above on these two additional datasets. For the BEAST analyses, all trees were summarized in a maximum clade credibility tree with the software Treeannotator (part of the BEAST package), after removing the burn-in and sub-sampling one tree every 10 000 generations.

## RESULTS AND DISCUSSION

### Phylogenetic inference of *M. bovis* and *M. caprae* populations

The phylogenetic reconstruction of all *M. bovis* and *M. caprae* sequences obtained (n=3364) confirmed that these two ecotypes correspond to two monophyletic groups, despite infecting similar hosts [49, 50] (Fig. S3). The range of host species from which *M. bovis* was isolated is broad, confirming that *M. bovis* can cause infection in many different mammalian species (Table S1). Our collection of *M. caprae* included genomes from Japan (isolated in elephants from Borneo) [51], China (isolated in primates, Table S1) and Peru (host information unavailable, but possibly human [52]), suggesting that the host and geographic distribution of this ecotype ranges well beyond Southern- and Central Europe [53, 54]. One group of *M. caprae* genomes with origin in Germany contained a deletion of 36-38 kb, which encompasses the region of difference RD4 (Table S1). This is in agreement with a previous study reporting Alpine *M. caprae* isolates as RD4 deleted, using conventional RD typing [55]. For all *M. bovis* genomes, x we determined *in silico* clonal complexes Eu1, Eu2, Af1 and Af2 and spoligotypes, and mapped them on the phylogenetic tree and onto a world map (Fig. 1, Fig. S2-S3, Table S1). All previously described clonal complexes corresponded to monophyletic groups in our genome-based phylogeny (Fig. S3). The phylogenetic tree also revealed *M. bovis* representatives that did not fall into any of the previously described clonal complexes (n=175, 5.3%, Fig. 1, Fig. S2-S3, Table S1). These belonged to eight monophyletic clades with unknown classification and to a few singleton branches (Fig. S3). The tree topology showed a strong bias towards closely related strains, in particular among Eu1, which reflects the different sampling and WGS efforts in the different geographic regions (Fig. 1, Fig. S3). Closely related genomes inform the local epidemiology but not the middle/long term evolutionary history of the strains and were thus excluded from further analysis (Subsampling 1, see Methods).

**Figure 1.**
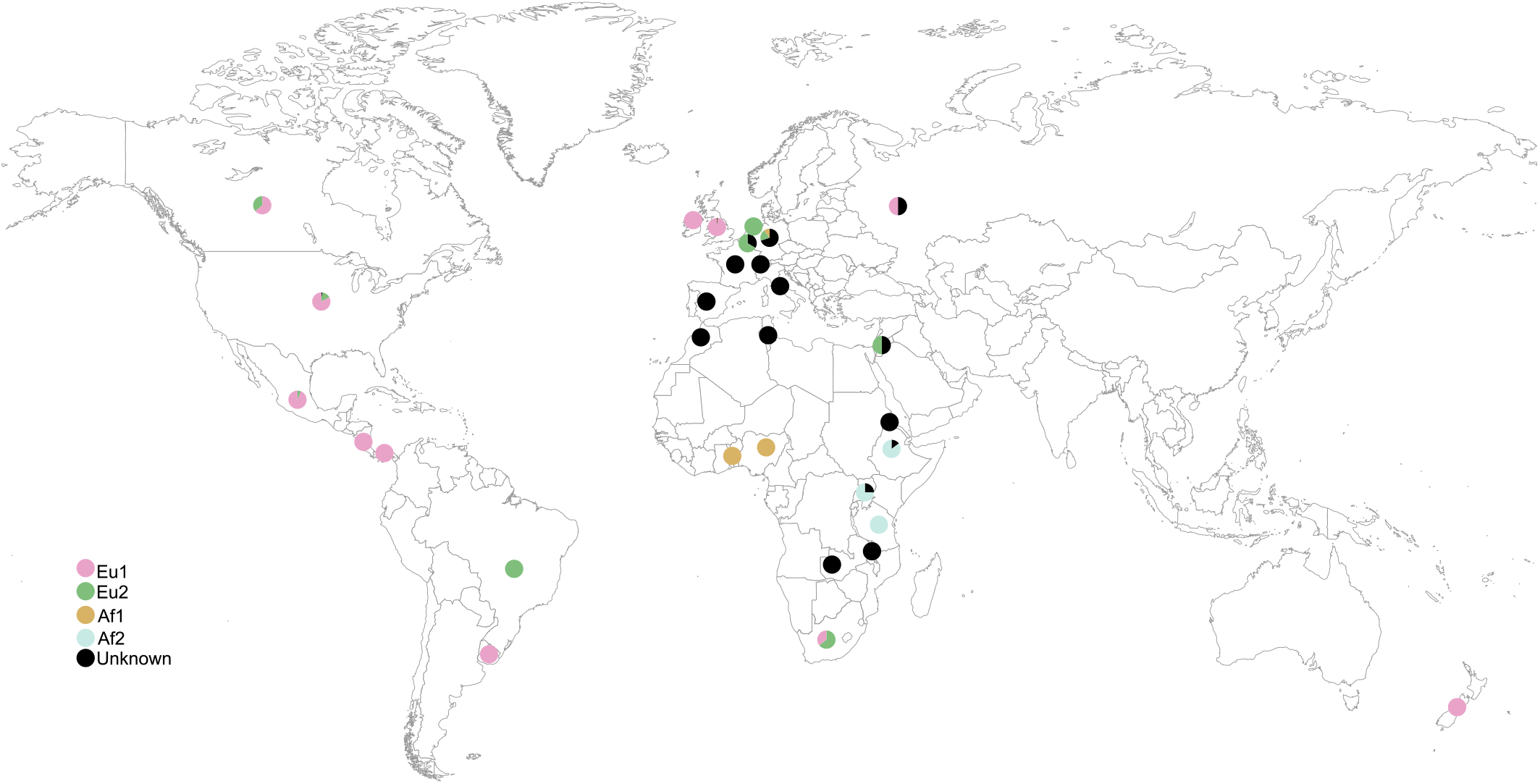
Geographic distribution of the *M. bovis* samples used in this study according to isolation country. The circles correspond to pie charts and are coloured according to clonal complexes.

**Figure 2.**
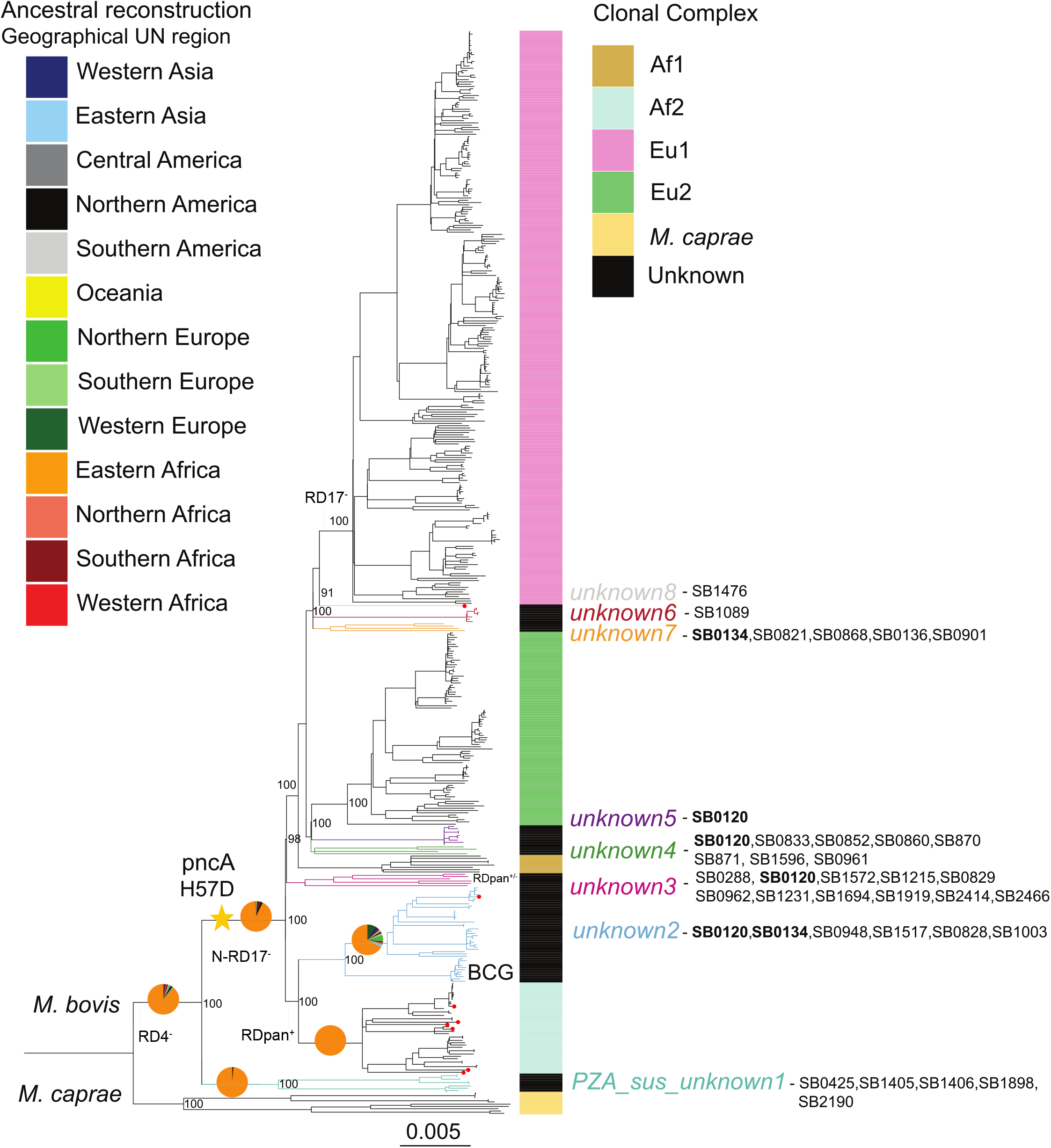
Maximum likelihood phylogeny of 476 of the 3364 genomes included in this study (redundant genomes were removed), and inferred from 22 492 variable positions. The scale bar indicates the number of substitutions per polymorphic site. The phylogeny is rooted on a *M. tuberculosis* Lineage 6 genome from Ghana (not shown) and bootstrap values are shown for the most important splits. The coloured bars on the side of the phylogeny show the different clonal complexes. Other “unknown” monophyletic clades are coloured in black and additionally the branches of the eight clades are coloured to show their phylogenetic position more precisely. The pie charts mapped on the tree represent the summary posterior probabilities (from 100 runs) of the reconstructed ancestral geographic states and are coloured according to geographical UN region. Inferred spoligotype patterns from WGS described in *M. bovis* spoligotype database [30] are indicated for the unknown clades. The red circles at the tips correspond to the eight newly sequenced genomes. Regions of difference (RD) as in [31] are indicated; superscript + and - refers to presence of the region or its deletion, respectively.

Two deep divergence events in *M. bovis* populations were notorious: one giving rise to an unclassified lineage we named *M. bovis PZA_sus_unknown1* (RD4 deleted as other *M. bovis*), which included five samples from Uganda (isolated from *Bos taurus* cattle, Table S1), three from Malawi (isolated from humans, Table S1) and one isolated from an antelope in Germany (Table S1). These *M. bovis* isolates lacked the *PncA* H57D mutation that is responsible for the intrinsic pyrazinamide resistance of canonical *M. bovis* as reported previously [56] (Fig. 2). This clade, retained the region of difference N-RD17, unlike all remaining *M. bovis*, and is probably related to a group of strains previously isolated from cattle in Tanzania and reported as “ancestral” by [31] (Fig. 2). In agreement with that, our *PZA_sus_unknown1* clade had other deletions reported as specific to these Tanzanian isolates; a larger deletion encompassing RDpan (RDbovis(a)_∆pan) and RDbovis(a)_kdp [31]. The second deep branching lineage included all other *M. bovis* strains descendent from an ancestor that acquired the *PncA* H57D mutation and therefore encompasses all previously described clonal complexes [8, 14], as well as the other previously unclassified clades we describe here.

From the *M. bovis* PZA resistant ancestor strains, two main splits occurred; one split led to the ancestor of Af2 and its previously unclassified sister clade which we called *unknown2* and which contains the BCG vaccine strains (Fig. 2). *M. bovis* strains with spoligotyping patterns similar to BCG have previously been referred to as “BCG-like”. However, our genome-based phylogeny shows that BCG-like spoligotyping patterns are present in several clades and have thus little discriminatory power [10] (Fig. 2, Table S4). In Af2 and *unknown2* the region of difference RDpan is present [31] (Fig. 2). Otherwise, RDpan is deleted in all other *M. bovis* except for the *unknown3* clade, in which it is polymorphic (Fig. 2). The other split led to the ancestor, from which all remaining *M. bovis* strains evolved, i.e. Af1, Eu2 and Eu1 as well as other groups (Fig. 2). Interestingly, Af1 does not share a MRCA with Af2 but with Eu1 and Eu2 as well as with another unclassified group, which we called *unknown3* (Fig. 2). Clonal complexes Eu1 and Eu2 share a MRCA together with five other *unknown* clades (*unknown 4* – *unknown 8*). Eu2 is more closely related to clades *unknown4* and *5*, than to Eu1 (Fig. 2). Eu1 in turn shares a common ancestor with three other clades *unknown6*, *7* and *8* (Fig. 2).

### The temporal and geographic origin of *M. bovis*

Our reconstruction of ancestral geographical ranges points to East Africa as the most likely origin for the ancestor of all *M. bovis* (Fig. 2, Fig. S4). This is supported by the fact that the basal clade *M. bovis* - *PZA_sus_unknown1* has an exclusively East African distribution and is pyrazinamide susceptible. Pyrazinamide susceptibility in *M. bovis* is probably an ancestral character given that all other lineages of the MTBC are pyrazinamide susceptible.

Alternatively, the ancestral *M. bovis* pyrazinamide susceptible populations could have had a much broader geographic distribution, which later became restricted to East Africa. For *M. caprae*, the sampling was too small and biased (Table S1) and no conclusions can be confidently drawn. We performed tip-dating calibration using the isolation dates of the strains with both Bayesian methods and LSD (see methods). Both the tip-to-root regression and the randomization tests performed indicated a temporal signal in the data (Fig. S5). We estimated a clock rate of between 6.66×10^−8^ and 1.26×10^−7^ for the BEAST analyses (Table S3), and between 6.10×10^−8^ and 8.29×10^−8^ for the LSD analysis. These results are in line with the results of previous studies [15, 57]. The common ancestor of *M. bovis* was estimated to have evolved between the years 256 and 1125 AD by the Bayesian analysis (cumulative range of the 95% Highest Posterior Density (HPD) of all nine BEAST analyses) and in the year 388 AD by LSD (Fig. 3, TreeS1-S10). Together, these estimates suggest that *M. bovis* has emerged in East Africa sometime during the period spanning the 3^rd^ to the 12^th^ century AD (Fig. 3). However, the credibility intervals of the different analysis spanned several centuries. This was due to the intrinsic uncertainty of dating ancestral nodes when the clock calibration is based on the sampling time of recently sampled tips (all strains considered in this study have been sampled in the last 40 years). Moreover, by relying exclusively on recent calibration points to date older nodes, we ignored the issue of the time-dependency of the estimated clock rates [58]. According to the time-dependency hypothesis, evolutionary rate estimates depend on the age of the calibration points, with older calibration points resulting in lower rates. In MTBC, this topic was discussed at length in other publications, and the available data to date does not allow to confidently accept or reject that hypothesis [57, 59–62]. Our analysis assumes that rates of evolution do not depend on the age of the calibration points. Therefore, we potentially underestimate the age of the older nodes of the tree. Molecular archaeological evidence suggests indeed that our dating analyses possibly underestimate the age of the MRCA of *M. bovis*. In particular, 2000 years old *M. bovis* DNA reported as having the RD4 and RD17 deletions, was found in human remains in Siberia [63]. The region of difference RD17 corresponds to RDEu1 [13], and defines the clonal complex Eu1 of *M. bovis*, which we estimate has evolved between the years 1236 and 1603 AD (Fig. 3).

**Figure 3.**
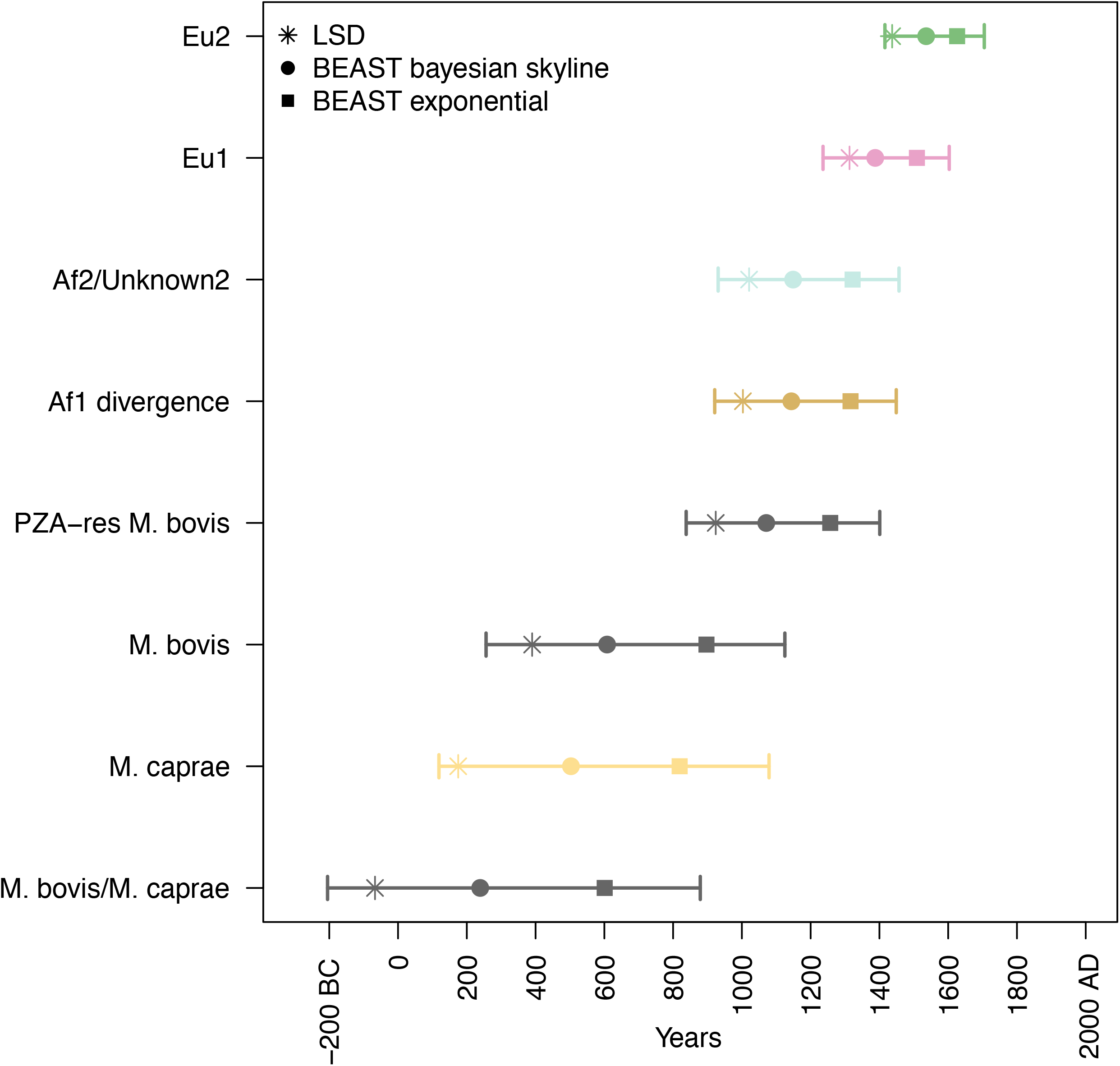
The inferred age of main monophyletic clades according to LSD and BEAST dating analyses. For BEAST we report the results of the two analyses that resulted in the lowest clock rate (subset1 Bayesian Skyline) and in the highest clock rate (subset3 exponential population growth). The confidence intervals reported correspond to the merged HPD interval of the two BEAST analyses mentioned above. The BEAST analysis was based on 300 genomes and the LSD analysis was based on 2058 genomes (see methods section for subsampling strategy). Only one genome from the Af1 clonal complex was included in the dating analyses and therefore the dates reported correspond to the node where Af1 diverged.

As discussed elsewhere [57], both tip-dating and the analysis of ancient DNA have potential pitfalls, and these discrepancies cannot be reconciled without additional data. Nevertheless, the tip-dating calibration provided accurate results for the emergency of BCG strains and for the introduction of *M. bovis* to New Zealand [64, 65](TreeS1-S10), indicating that the method can reliably infer divergence times at least for events occurred in the last 200 years.

### Insights into the detailed population structure of *M. bovis* around the world

Understanding the evolutionary history of the *M. bovis* populations requires understanding their geographic distribution at a continental scale. Our WGS data set has limited geographical resolution due to the biased sampling of certain regions of the world, and to the partial unavailability of associated metadata such as the origin of foreign-born TB patients from Western countries. To get more insights into the geographical ranges of the different *M. bovis* clades, we used the spoligotype patterns inferred from the WGS data and searched for references describing the prevalence of those in different regions of the world (Table S4). Patterns SB0120 and SB0134, known as “BCG-like” and reported to be relatively prevalent [9], as well as SB0944, are phylogenetically uninformative; SB0120 is present in several clades, and SB0134 and SB0944 have evolved independently in two different *M. bovis* populations (Fig. 2, Table S4).

Our results suggest that the sister clade of all contemporary pyrazinamide resistant *M. bovis*, *PZA_sus_unknown1*, is restricted to East Africa. The same holds true for Af2, which is in accordance with previous reports [8, 12]. Our findings further suggest that the geographical distribution of the Af2 sister clade *unknown2* includes East Africa (Eritrea, Ethiopia), but also Southern Europe (Spain and France). Informative spoligotypes of the *unknown2* clade show that it also circulates in North Africa (Fig. 2, Table S4). Of note, the original strain, from which all BCG vaccine strains were derived, was isolated in France [66]. Our inferences suggest that a common ancestor of Af2 and *unknown2* evolved in East Africa, and while Af2 remained geographically restricted, its sister clade *unknown2* has subsequently dispersed (Fig. 2).

All remaining *M. bovis* descended from a common ancestor, for which the geographical origin was impossible to infer reliably with our data. However, the tree topology showed that from this ancestor several clades have evolved which are important causes of bovine TB today in different regions of the world (i.e., the clonal complexes Eu1, Eu2 and Af1; Fig.2).

The most basal clade within this group is *unknown3*, which contained 25 genomes mostly isolated from humans (Table S1). The *in silico* derived spoligotypes suggest that the geographical spread of *unknown3* ranges from Western Asia to Eastern Europe, but also includes East Africa (Fig. 2, Table S4). The next split in our phylogeny corresponds to Af1, which has been characterized extensively using the deletion RDAf1 and spoligotyping, and shown to be most prevalent in countries from West- and Central Africa [11]. Here, we could only compile nine Af1 genomes, of which five originated in Ghana [67], and the remaining had either a European or an unknown origin. The small diversity of Af1 spoligotypes found in our WGS dataset [11] indicates strong undersampling (Fig. 2, Table S4). Nevertheless, it was possible to estimate the divergence of the Af1 clade from the remaining *M. bovis* to a period ranging from the year 921 to 1449 AD (Fig. 3), making it unlikely that Af1 was originally brought to West Africa by Europeans [68].

The next split comprises clades *unknown4*, *unknown5* and Eu2. Clade *unknown4* was composed of 33 genomes with little geographic information and for which the most common spoligotyping pattern was the uninformative SB0120 (n=19). Additional *unknown4* spoligotypes indicate that strains belonging to this clade circulate in Southern Europe, Northern and Eastern Africa (Fig. 2, Table S4), supporting dispersion events between Africa and Southern Europe. Clade *unknown5* comprised only nine genomes isolated mostly from Zambian cattle. Its corresponding spoligotype is also SB0120, limiting further geographical inferences.

In contrast to the strains from clades *unknown4* and *unknown5*, among the 323 Eu2 genomes, no genomes of East African origin were found, and Africa was only represented by nine South African genomes [69]. By far, most Eu2 were isolated in the Americas. Previous studies have shown that Eu2 dominates in Southern Europe, particularly in the Iberian Peninsula [14], thus possibly the source of Eu2 in the Americas. There were no representatives of Eu2 from the Iberian Peninsula in our dataset. However, our molecular dating analysis revealed that the common ancestor of Eu2 evolved during the period 1416 to 1705 AD (Fig. 3), which would be compatible with an introduction from Europe into the Americas.

Clonal complex Eu1, *unknown6*, *unknow7* and *unknown8*, form a sister group to the previously described. Eu1 has previously been characterized based on the RDEu1 deletion and spoligotyping, showing that it is highly prevalent in regions of the world that were former trading partners of the UK [8, 13]. That geographic range is well represented in our dataset, including many genomes from the UK (n=215) and Ireland (n=45) (Table S1). The latter were very closely related, suggesting that there was probably fixation of just a few genotypes in this region as previously proposed [8]. In contrast, most branching events within Eu1 correspond to *M. bovis* isolated in North- and Central America as well as New Zealand, resulting from the expansion of clonal families not seen in the British Islands. Consequently, most of the genetic diversity of Eu1 exists outside of its putative region of origin. Our molecular dating is compatible with this view, indicating that the ancestor of Eu1 is likely to have emerged between the years 1236 to 1603 AD (Fig. 3), with several Eu1 sub-clades evolving in the last 200-300 years (TreeS1-S10).

The closest relative to Eu1 is a genome from Ethiopia (*unknown8*) with the spoligotyping pattern SB1476, commonly found in Ethiopia [12]. *Unknown6* comprised seven genomes from North America (Fig. 2, Table S1, Table S4), whereas *unknown7* included eight genomes, four of which were isolated in Western Europe and another four without country of origin available. Spoligotyping patterns indicate that identical strains are common in Southern Europe, Northern and Eastern Africa expanding the geographic range of *unknown7*.

## CONCLUSIONS AND IMPLICATIONS

We screened the public repositories and compiled 3364 genome sequences of *M*. *bovis* and *M. caprae* from 35 countries. Despite the biased geographic distribution of our samples, our results provide novel insights into the phylogeography of *M. bovis* and *M. caprae.* Our whole-genome based phylogeny showed that although certain spoligotypes are associated with specific monophyletic groups, prevalent patterns such as the so-called “BCG-like” should not be used to infer phylogenetic relatedness. Moreover, our data extend the previously known phylogenetic diversity of *M. bovis* by eight previously uncharacterized clades in addition to the four clonal complexes described previously. Among those, *Pza_sus*_*unknown 1* shares a common ancestor with the rest of *M. bovis*, has an exclusively East African distribution and does not share the PncA mutation H57D, conferring intrinsic resistance to PZA.

Our further inferences suggest that *M. bovis* evolved in East Africa. The evolutionary success of *M. bovis* is linked to the fact that it can infect and transmit very efficiently in cattle. Cattle have been domesticated twice independently; once in the Near East (*Bos taurus*) and once in the Indus Valley (*Bos indicus*) approximately 10 000 year ago, and both were introduced to Africa at different time points and various locations, subsequently interbreeding with local wild species [70]. Whereas *B. taurus* was introduced probably during the 6 millennium BC possibly through Egypt, *B. indicus* was most likely introduced twice, first during the second millennium BC and later during the Islamic conquests [71]*. M. bovis* could have emerged after the introduction of cattle, benefiting from the development of African pastoralism and expanding within the continent. The timing of these events is difficult to estimate; the initial introductions of cattle predate by several thousands of years our inferred temporal origin of *M. bovis*. But as discussed, our estimates are possibly affected by the current uncertainty in dating deeper evolutionary events within the MTBC. Alternatively, *M. bovis* could have emerged in the Near East and been introduced to Africa together with cattle. We cannot test this hypothesis formally, as the Near East is poorly represented in our dataset. However, this scenario is difficult to reconcile with the restricted East African distribution of the *Pza_sus_unknown_1* clade, as the Near East was also the origin of taurine cattle both in Europe and Asia, where no clade retaining the ancestral characteristic of pyrazinamide susceptibility was found.

While some *M. bovis* groups remained restricted to East Africa, others have dispersed to different parts of the world. The contemporary geographic distribution of *M. bovis* clades suggest that East- and North Africa, Southern Europe and Western Asia have played an important role in shaping the population structure of these pathogens. However, these regions were not well represented in our dataset. Thus, more *M. bovis* genomes from these regions are necessary to generate better insights, particularly given the central role of these regions in the history of cattle domestication [72]. From a more applied perspective, our work provides a global phylogenetic framework that can be further exploited to find better molecular markers for studying *M. bovis* in settings where genome sequencing cannot be easily implemented.

## ACKNOWLEDGEMENTS

Calculations were performed at sciCORE (http://scicore.unibas.ch/) scientific computing core facility at University of Basel. This work was supported by the Swiss National Science Foundation (grants 310030_166687, 310030_188888, IZRJZ3_164171, IZLSZ3_170834 and CRSII5_177163), the European Research Council (309540-EVODRTB) and SystemsX.ch.

## Supplementary Figures

**Figure S1** – Flow chart showing the selection of genomes.

**Figure S2** - Geographic distribution of the *M. bovis* samples with unknown classification used in this study according to isolation country.

**Figure S3** – Maximum likelihood phylogeny of all 3364 genomes, based on 45 981 variable positions. The scale bar indicates the number of substitutions per polymorphic site. The phylogeny is rooted on a *M. tuberculosis* Lineage 6 genome from Ghana. The outer ring indicates the geographical region from which the strains were isolated. The four clonal complexes are highlighted on the tree. Branches corresponding to BCG genomes are coloured in grey and the *PncA* mutation H57D is indicated by a yellow star.

**Figure S4** – Phylogeographic reconstruction of *M. bovis* and *M. caprae*, inferred from 392 genomes. Thirteen UN-defined geographic regions were assigned to the discrete character geographic origin, and mapped onto the phylogeny. Pie charts at internal nodes represent the summary posterior probabilities (from 100 runs) of the reconstructed ancestral geographic states and are coloured according to geographical UN region.

**Figure S5** – A) Tip-to-root regression and B) Date randomization tests (DTR). The confidence interval of the clock rate estimate for the observed data does not overlap with the confidence intervals of the clock rate estimates obtained from the randomized sets.

## Supplemental Tables

**Table S1 –** List of genomes included in this study along with metadata used for the analyses.

**Table S2 –** Comparison of models for discrete character evolution using likelihood ratio tests.

**Table S3 –** Results of all BEAST analyses.

**Table S4 –** Spoligotype patterns determined *in silico* for different clonal complex groups with reference to other studies.

## Supplementary files

**TreeS1 –S10** Ten time-calibrated trees resulted from the molecular clock analyses. The file names indicate the software used (LSD or BEAST), the subsample, and the coalescent population prior (BSP: Bayesian Skyline; exponential: exponential population growth; constant: constant population size). Tip labels are present in Table S1. Ages in years before present can be visualized as well as the 95% High Posterior Density (HPD) (BEAST trees) using FigTree [73].

