## Supplementary figures and images for "An African origin for *Mycobacterium bovis*"

### FigS1

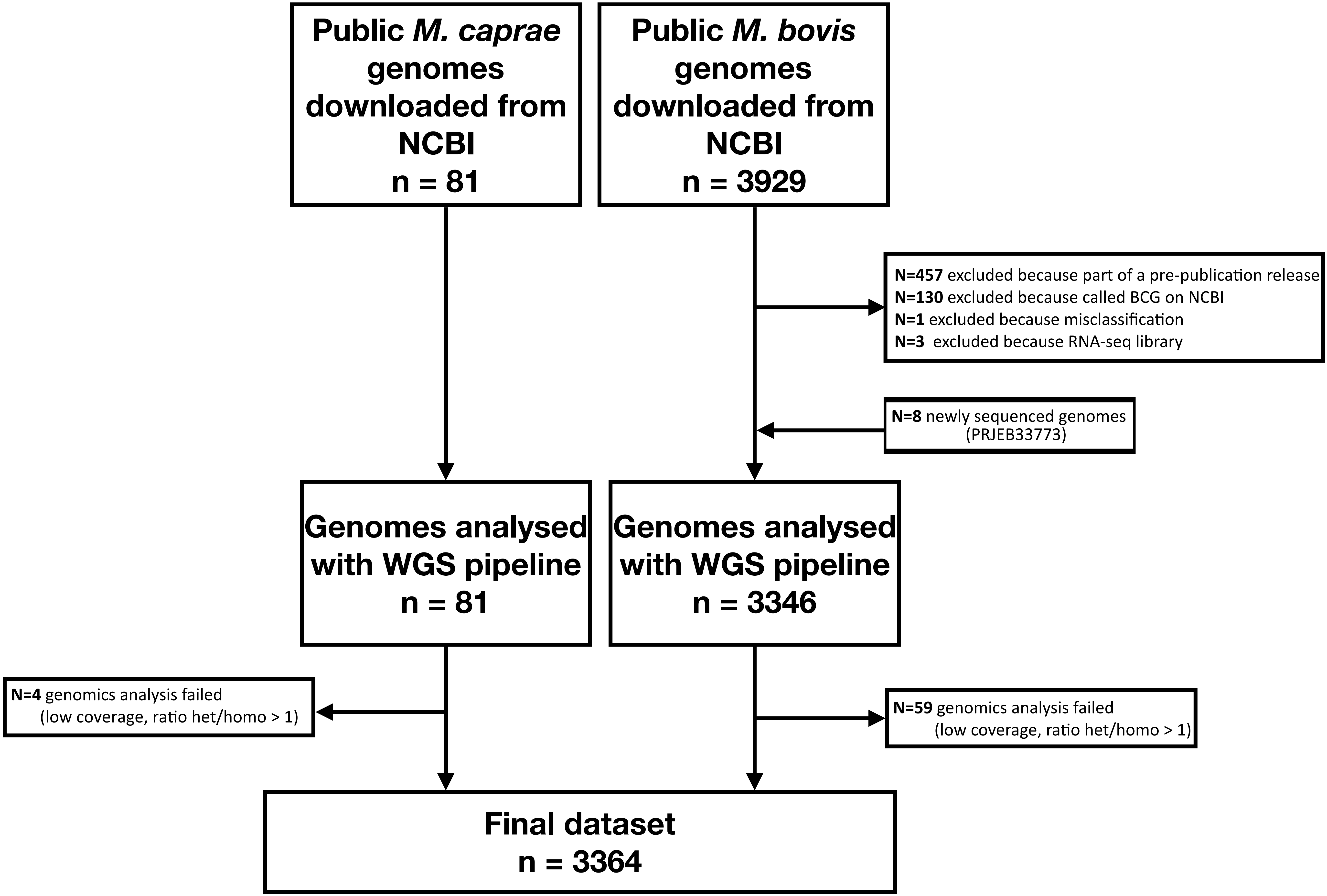

### FigS2

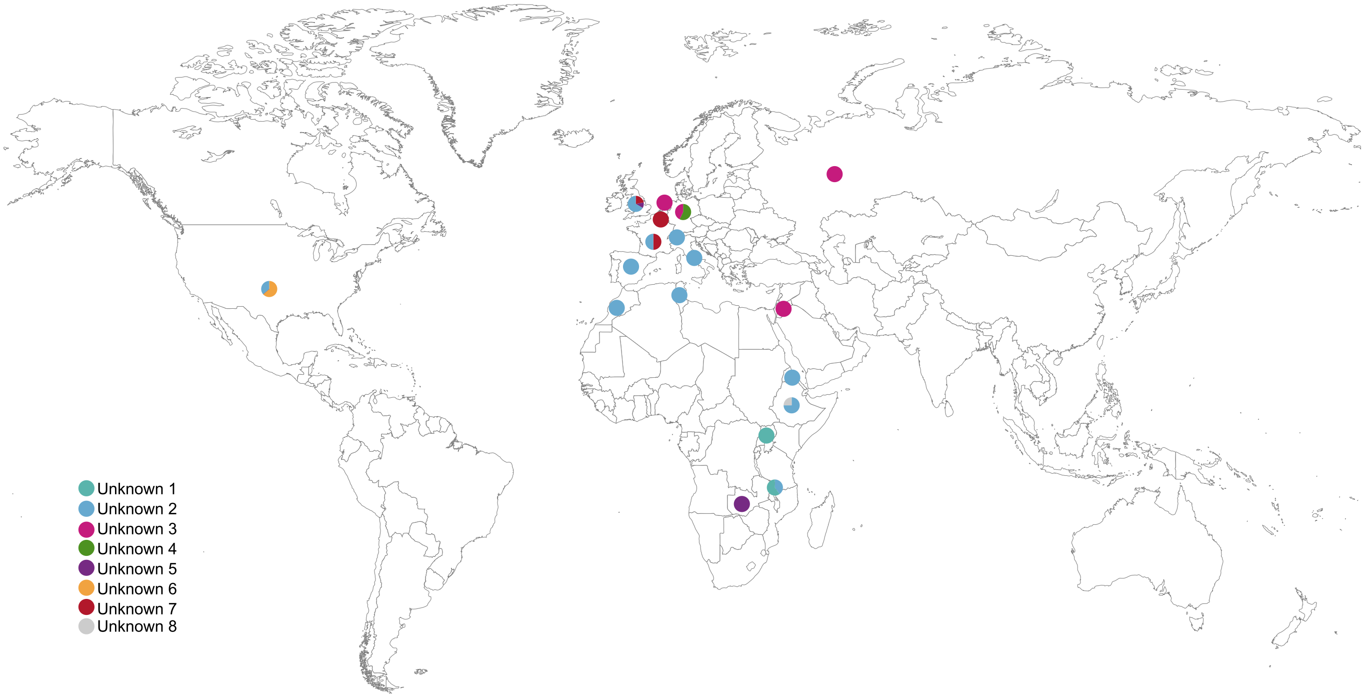

### FigS3

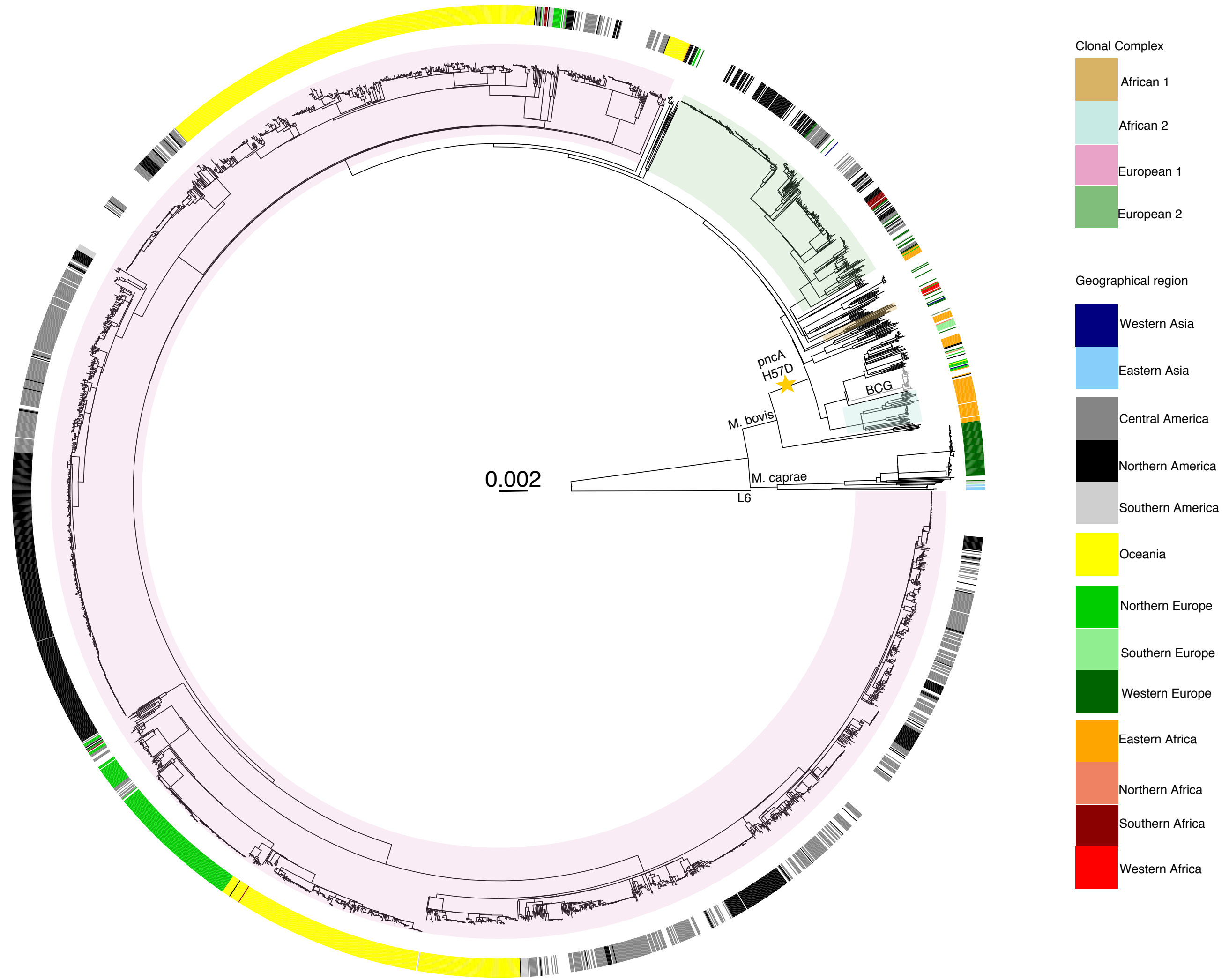

### FigS4

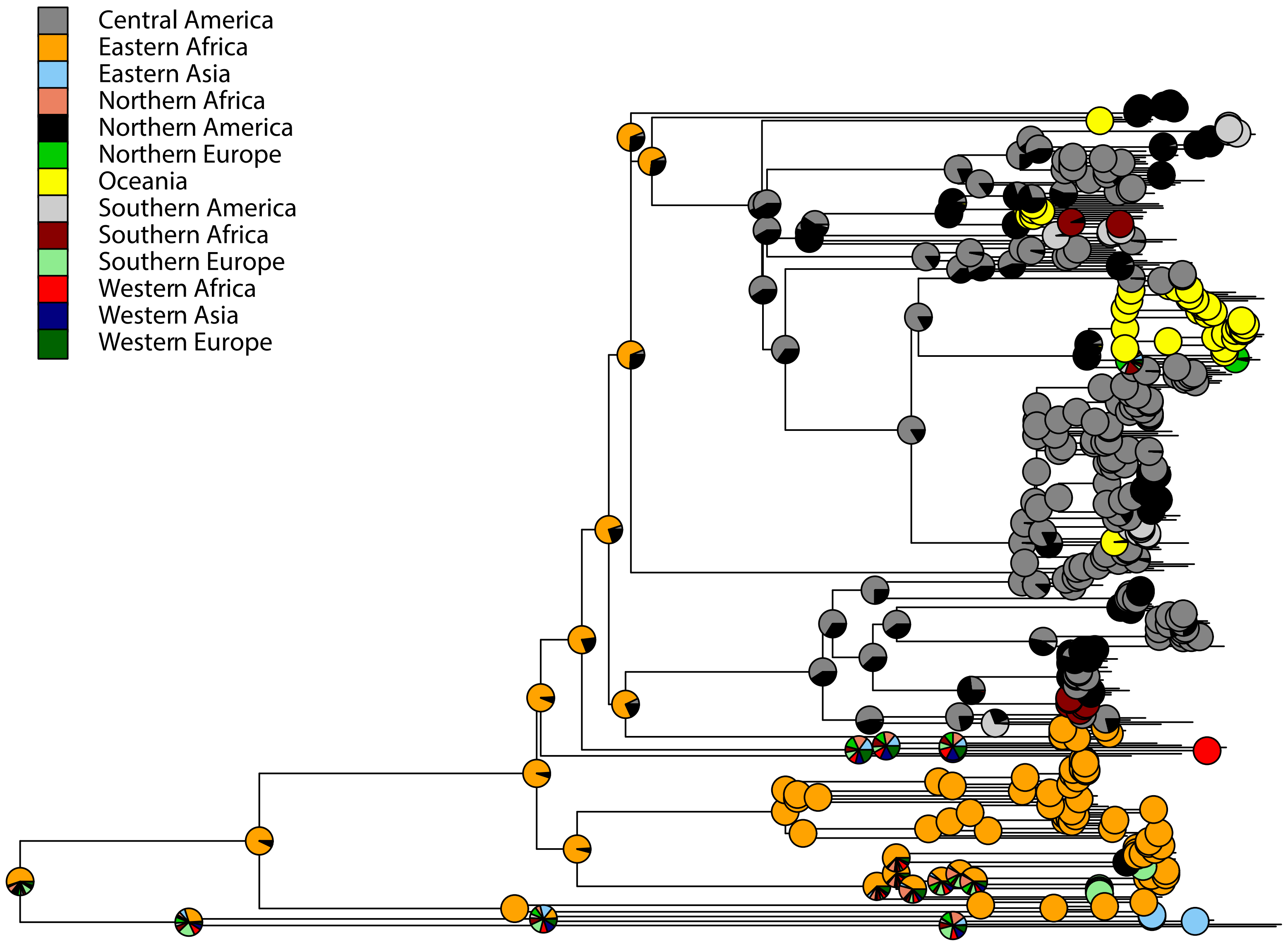

### FigS5A

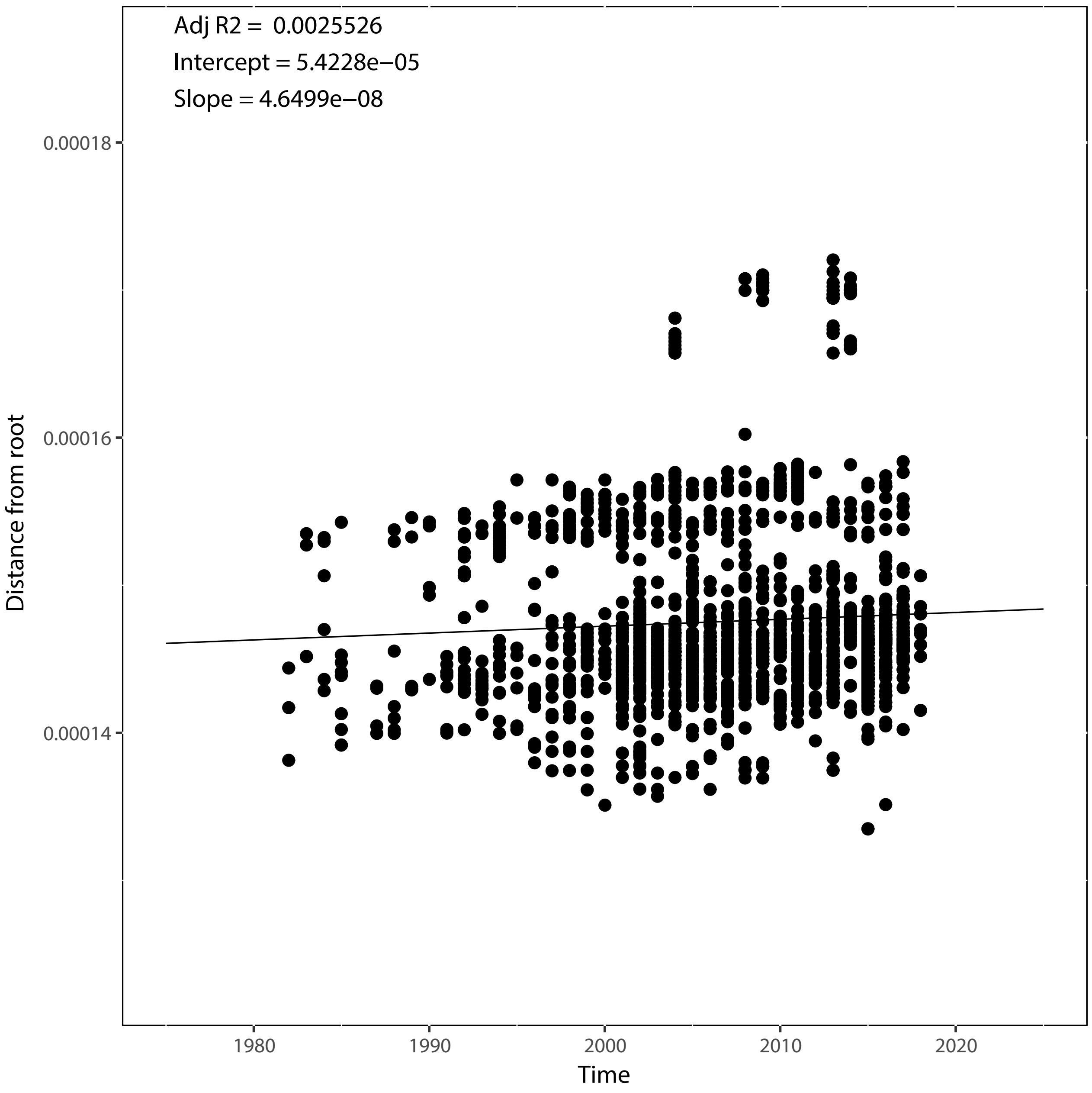

### FigS5B

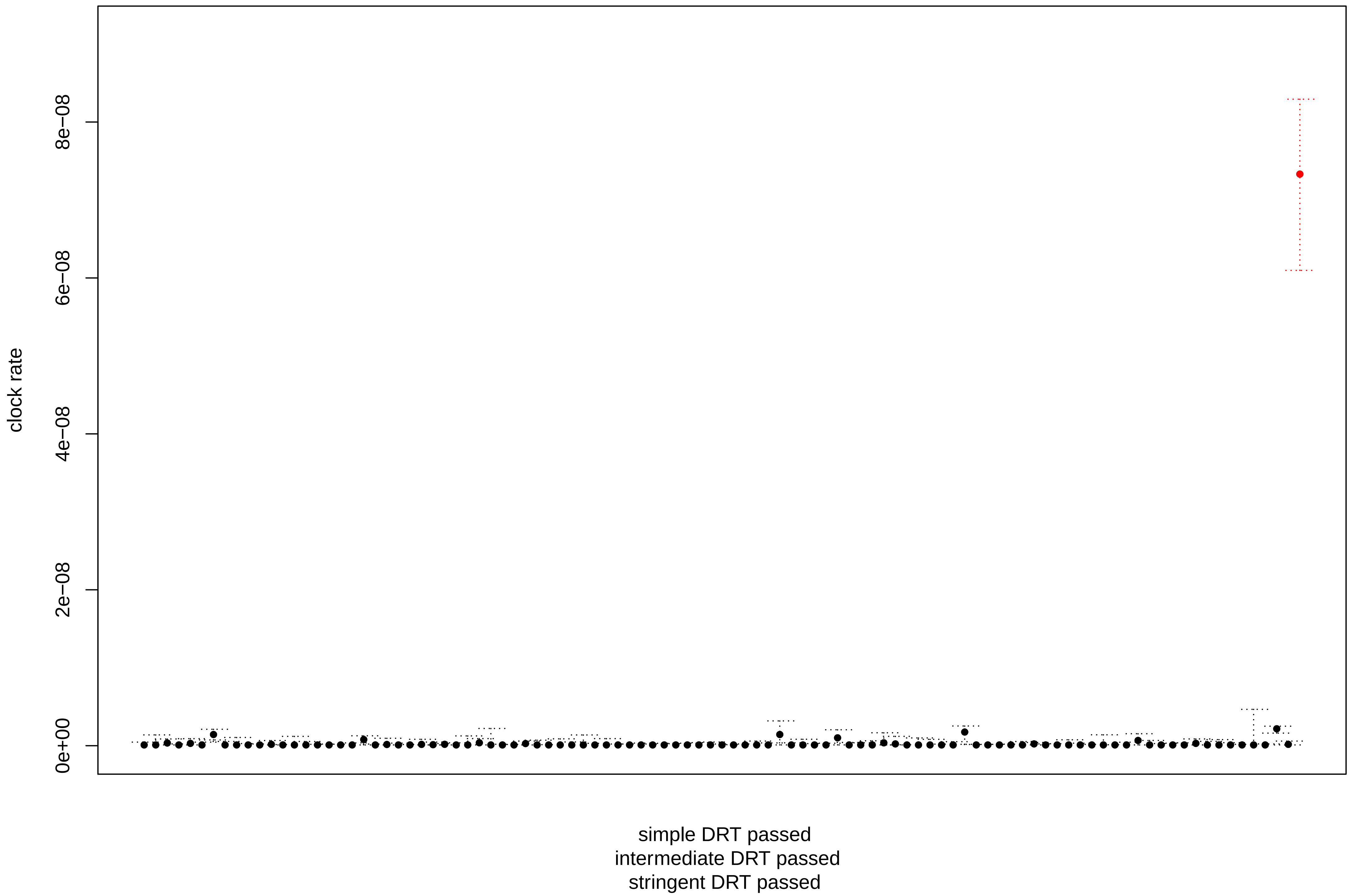
